# A Pose-Informed De-Noising Diffusion Model for Adult Naturalistic EEG Signals

**DOI:** 10.1101/2023.12.08.567146

**Authors:** Angshuk Dutta, Marcel Hirt, Lorena Santamaria, Stanimira Georgieva, Christian Gerloff, Boyang Li, Victoria Leong

**Affiliations:** School of Social Sciences, Nanyang Technological University, Singapore; Lehrstuhl II für Mathematik, RWTH Aachen, Germany; School of Computer Science and Engineering, Nanyang Technological University, Singapore; Dept of Pediatrics, University of Cambridge, United Kingdom

**Keywords:** EEG, Diffusion Models, Generative Modelling

## Abstract

Artifact contamination in EEG (electroencephalogram) signals is a significant problem, especially in naturalistic settings where participants can move freely. This contamination stems from various sources like eye movements, muscle activity, sweat, and electrical interference, whose effects differ greatly from each other. Traditional denoising methods, such as Independent Component Analysis, are limited because they assume a linear relationship between the source of the artifacts and the EEG signals, and often require the dominance of one noise source over others. Moreover, these methods need expert knowledge in EEG analysis and lack an objective standard for evaluation.

To overcome these challenges, we propose two innovations: Firstly, we introduce the use of “video-estimated” pose coordinates – the x and y positions of different body points (like wrists, eyes, and ankles) – to assist in the EEG denoising process. Secondly, we present a denoising diffusion model, EEG-DDM, that utilizes both the contaminated EEG signals and these pose coordinates to effectively denoise the EEG. Our findings show that incorporating keypoints (pose coordinates) improves denoising performance and helps maintain cross-spatial dependencies in the data. Additionally, we enhance human interpretability of the process by displaying saliency maps generated by our model, which explain the contributions of these keypoints in the denoising process.

## I. Introduction

The Electroencephalogram (EEG) has long been as a preferred technique for neural data acquisition in adult populations, primarily due to its superior temporal resolution and the non-invasive approach [1]. However, it also brings several challenges such as high sensitivity to noise, which can mask the underlying neural signals [2]. Examples of said noise include ocular [3], muscular [4], electrocardiogram (ECG) [2] and line noise. While line noise can be easily removed [5], ocular and muscular artifacts pose a significant challenge due to their complex broadband effects on the signal.

In naturalistic experimental paradigms, EEG movement artifacts are amplified. Thus, removal of such artifacts is of paramount importance and several methods have been devised to do so. For adult populations, various classes of methods exist such as wavelet decomposition, blind source separation (BSS), artifact subspace reconstruction (ASR), empirical mode decomposition, canonical correlation analysis and so on. The one considered most reliable is independent component analysis (ICA) [6], which is a BSS method. ICA decomposes the EEG signal into several independent components which can be classified as originating from the brain or artifactual. The removal of these artifactual ICs thus recovers the denoised signal. ASR [7] denoises EEG in a particular window at a time whose size is a hyperparameter. ASR detects segments with the highest variance and aims to remove these segments. To do so however, ASR need to be calibrated using a reference signal or clean EEG. Another popular method introduced is wavelet decomposition, where — similar to ICA — the signal can be decomposed into several wavelet coefficients and identifying a suitable threshold can help remove noisy coefficients.

However, these methods still have certain challenges. For instance, a formal proof of the limitation of ICA provided by [8], suggests that the success of ICA is contingent on the distribution of the noise variance being uniform and the data being full rank which may not always be the case due to the spectral properties of the different artifacts. In the case of wavelet decomposition, the quality of artifact removal depends on the choice of the mother wavelet and threshold creation which requires expertise and may be time consuming. In both cases, we can generalise the limitiations to the choice of the basis functions (ICs for ICA and wavelet coefficients for wavelet decomposition) that these methods project the data to.

Several deep learning approaches such as denoising EEG signals with Generative Adversarial Networks [9], removing ocular and myogenic artifacts with deep learning [10] have been developed which overcome the limitations of traditional EEG denoising methods by learning flexible basis functions from the data itself. However, deep learning methods, being highly non-linear, are not interpretable.

Thus to aid in solving the interpretability problem and improving model performance, we introduce pose coordinates which are estimated from video recordings of the participants using machine learning algorithms like OpenPose [11]. These keypoints consist of a multivariate time series of several key pose features on the body such as eyes, nose, wrists, hips and similar others as depicted in fig 1. We also introduce a keypoint saliency model to explain the contribution of these keypoints in the denoising process. We hypothesise that EEG movement related artifacts are directly related to the distance and speed of change in the position of the relevant body part(s) that generate motion. For example, a large variation in the x and y coordinates of the eyes, nose, neck and ears may indicate head motion artifact at that corresponding time point in the EEG. Finally, we introduce a reverse denoising diffusion model (EEG-DDM) which utilises these keypoints along with the artifact corrupted EEG to denoise the EEG signals.

**Fig. 1:**
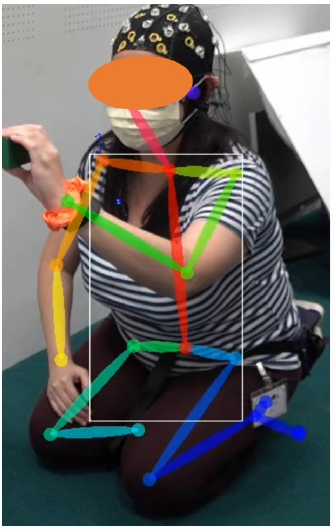
A subset of the pose coordinates of body keypoints collected from videos. Written informed consent was obtained for the publication of this image

## II. Materials and Methods

### A. Dataset

The dataset comprised of EEG recordings from 35 healthy adult women with a mean age of 32.17 years and standard deviation of 4.12 years. in two distinct tasks with their infants: the A-not-B task [12] and the Sequential Touching Task (STT) [13]. In the A-not-B task, mothers were seated across their infants with a tray containing two covered bowls. The mother placed a toy in one of these bowls and used verbal and facial cues to encourage the infant to identify the correct bowl.The STT was divided into three stages pre-play, demonstration and post play. In this paper we focused on the demonstration phase since the mother actively moves and interacts with her infant while demonstrating the toy’s features leading to more artifacts while other phases involve the mother being seated behind the child with limited interaction. Although EEG data was also collected from the infant for both tasks, only the mother’s EEG will be reported in this paper. Ethical approval for this research was obtained from the Nanyang Technological University IRB (IRB-2021-808).

### B. EEG data acquisition

EEG was recorded using a 32-channel cap (30 channel EEG and 2 channel ECG) and was amplified using a LiveAmp from Brain Products. The EEG data was recorded using the Brain Vision recorder software at a 500 Hz sampling rate and referenced to the Cz electrode. The impedance was under 20kΩ. The adult EEG LiveAmp and the video recordings were synchronised using a push button trigger system which delivered a trigger marker to the EEG and simultaneously a LED light for all the video cameras that were used to monitor the task.

### C. EEG prepropocessing

For the ground truth, the EEG was manually preprocessed using ICA. The EEG signal was high-pass filtered at 0.5 Hz and low-pass filtered at 40 Hz to obtain good quality ICA components. Bad channels were identified visually and were removed prior to average referencing. Cz was recovered in the average referencing. Large multi-channel artifacts were annotated and excluded from the ICA algorithm (runica option in the EEGLAB [14] plugin for MATLAB). The IC components were visually inspected and artifact ICs were removed. The recovered EEG was then used as the ground truth to compare against the proposed model performance.

### D. Video recordings and Keypoint Extraction

Videos were collected for the task using 3 Sony FDR-AX700 4K resolution cameras with a frame rate of 25 frames per second. Each of these cameras were positioned at three distinct locations. One camera was placed to only record the mother thus excluding the infant from the field of view and vice versa. The final camera was placed to the side capturing a sideways view of both the mother and the infant. The mother camera was used to extract the adult keypoints by passing the adult viewpoint video to the OpenPose [11] pipeline. These motion parameters estimated contain 2D coordinates (x and y) of the 25 body features and thus result in a total of 50 features per frame.

### E. EEG Artifact model

Consider *x*_clean_ ∼ *p*(*x*_clean_) to be the clean EEG sampled from the unknown probability distribution *p*(*x*_clean_) and *x*_clean_ ∈**R**^*C× 𝒯*^ where *C* is the number of channels and 𝒯 is the number of samples in time respectively. Further, let *x*_artifact_ ∼ *p*(*x*_artifact_) denote the artifact contaminated EEG sampled from the distribution *p*(*x*_artifact_). Following the well known generative assumption that *x*_artifact_ can be modeled as

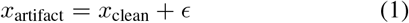

where *ϵ* represents the noise signal, we aim to directly learn a different decomposition similar to equation 1 but *ϵ* ∼ 𝒩 (0, *σ*(*x*_artifact_, *k*)^2^*I*) where *k* ∈ **R**^*C′× 𝒯 ′*^ denotes the multivariate keypoint times series consisting of 𝒯^*′*^ samples from *C*^*′*^ key body features as explained in section II-D. *ϵ* is now estimated as a function of the artifactual EEG and the pose coordinates. To estimate *ϵ* or rather *σ*(*x*_artifact_, *k*), we introduce the keypoint saliency model in section II-G and use *σ*(*x*_artifact_, *k*) in our generative diffusion model to obtain *x*_clean_ from *x*_artifact_ as discussed in section II-F

### F. Diffusion model framework

We suggest to model the artifact-corruption process through a diffusion model as compared to other generative models due to their superior generative performance [15] and stable training . More precisely, we assume for the forward diffusion process a VE-SDE (Variance Exploding Stochastic Differential Equation) parameterisation, cf. [16],

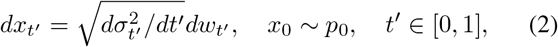

where *σ*_*t*_*′* is the variance at time *t*^*′*^ and *w* a Wiener process. In this framework *x*_0_ denotes the clean EEG samples and *p*_0_ denotes its distribution such that (2) describes the transition of a clean EEG sample into a noisy sample *x*_1_ ∼ *p*_1_. We assume that the observed EEG data *x*_artifact_ corresponds to *x*_*t*_ for some *t*≤ 1. Following a similar approach [17] developed for MR image denoising, we consider a standard variance schedule at time *t*^*′*^ as *σ*_*t*_*′* = *σ*_*min*_ (*σ*_*max*_*/σ*_*min*_)^*t*^*′* where *σ*_*max*_ and *σ*_*min*_ are hyperparameters. Given *x*_*t*_∼ *p*_*t*_ and our forward diffusion process, we can sample *x*_0_ which is the clean EEG by iteratively solving the reverse diffusion process by using numerical solvers described in [16]. See section 3.3 and 4 in [16] for model details. However, not knowing the diffusion time *t* may lead to longer sampling time or non convergence to the clean sample. Given our variance schedule, we can estimate *t* if the variance *σ*_*t*_ is known. However estimating the noise variance *σ*_*t*_ from *x*_*t*_ is a nontrivial task that we address by utilising the video estimated pose coordinates.

### G. EEG Noise Variance Estimation using Keypoints

As explained in section II-E, our proposed framework for denoising EEG requires the estimation of the diffusion time *t* at which the artifactual EEG *x*_*t*_ is generated. Thus, we introduce a keypoint saliency model which utilises the keypoints *k* and the artifactual EEG *x*_*t*_ as input to estimate the noise variance *σ*_*t*_ and in turn estimate the diffusion time *t*.

For data preprocessing, we introduce a series image 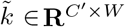 where *C*^*′*^ represents the number of body pose features and *W* represents the number of samples in time and *W <* 𝒯 ^*′*^ where 𝒯 ^*′*^ is the total number of samples in time and 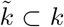. Furthermore, we augment 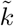 by adding Gaussian noise or applying gaussian blur to random segments. This form of data augmentation can help learn trend variations effectively as well as enhance local information and stabilise training. However, training the model using data augmented with this technique makes it sensitive to the noise variance added. To prevent such biases, we utilise a learnable mask M∈ [0, 1]^*C′×W*^ as introduced by [18] and transform 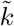 as follows:

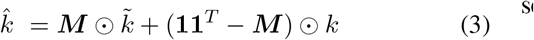

where **1** is a vector of ones and ⊙ is the Hadamard product.

For the model architecture, we utilise a Transformer Encoder with 6 encoder blocks whose key and query input variables are the augmented series image 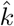and the artifact contaminated EEG *x*_*t*_ respectively. The variance *σ*_*t*_ estimated by the keypoint model is used to predict the noise *ϵ*_pred_ using the reparametrisation trick [19]. The model is then optimised using the loss function

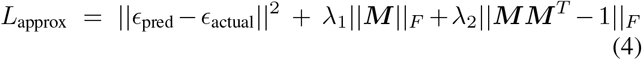

where the first loss term represents the reconstruction loss and *ϵ*_*actual*_ represent actual noise given by *x*_*t*_ − *x*_0_. *λ*_1_ and *λ*_2_ are hyperparameters. ||***M***|| _*F*_ represents the Frobenius norm of the mask M and prevents the mask from converging to **11**^*T*^ such that we obtain the contributions of the distinct keypoints. The final term ||***MM*** ^*T*^− 1 ||_*F*_ allows for spatial permutation as ***M*** approaches orthogonality and better model performance as demonstrated by [18]

### H. Generating Interpretable Keypoint Contribution

To explain the contributions of different body keypoints towards the estimation of the noise variance *σ*_*t*_, we generate saliency maps by making use of the keypoint saliency model. The saliency maps are represented by the mask ***M*** and are obtained by optimising only ***M*** while keeping the parameters of the trained saliency model fixed. Thus, the optimal mask ***M*** is obtained by minimising the following loss function

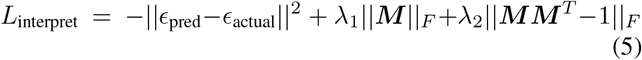

where *ϵ*_pred_ is the noise sampled from the keypoint saliency model and *ϵ*_actual_ is the noise predicted by the diffusion model. Note that the first term of equation (5) now has a negative sign as compared to equation (4) while the other two terms remain the same. Given that saliency model parameters are fixed, minimising the first term enables learning of a mask M to augment the input keypoint so as to worsen model performance. Thus the mask M represents the smallest destroying region (SDR) [20] and thus captures the important keypoints for the estimation of *σ*_*t*_

## III. Results

### Data Split and Model Training

The EEG and pose coordinate data from the N=35 participants was randomly split into 20 participants for training and 15 participants for testing. Furthermore, each EEG dataset was subdivided into 3s windows (1500 samples) and batched. Consequently the keypoints were also subdivided into 3s windows (75 samples) corresponding to the EEG. Both EEG-DDM and the keypoint saliency models were trained using a 19 fold cross validation strategy such that these models would not overfit on a specific subset of the participant dataset. All the results depicted below were obtained from executing the models on the 15 unseen datasets. For model comparisons IC-UNet, ICLabel and deep separator were considered to be the baseline models.

To demonstrate the efficacy of the keypoints we conduct two experiments to address the following questions:

- Does the score based model approximate the clean EEG better with the inclusion of pose coordinates from body keypoints?
- Does the addition of pose coordinates help preserve spatial covariance of the estimated denoised EEG?

#### A. Keypoints help Score Based Models to approximate Clean EEG

We test this hypothesis by utilising two different measures: relative root mean square error (RRMSE) and the correlation coefficient (CC). The relative root mean square error (RRMSE) is defined as the ratio of the mean of square root of residuals squared to the mean of observed values. For RRMSE, by definition, the lower the value the better. Further, the Correlation Coefficient (CC) is a similarity metric which is bounded between -1 and 1 with 1 being positively correlated and -1 being inversely correlated. In our case, the higher the correlation coefficient the better.

**Table 1:** Summary of the RRMSE and CC results depicted in fig 2

| Model | RRMSE(%) | CC |
| --- | --- | --- |
| Deep Separator | $55.357 \pm 2.530$ | $0.753 \pm 0.024$ |
| ICLabel | $43.307 \pm 2.848$ | $0.736 \pm 0.014$ |
| IC-UNet | $34.229 \pm 1.826$ | $0.809 \pm 0.018$ |
| EEG-DDM w/o keypoints | $22.281 \pm 1.278$ | $0.813 \pm 0.0156$ |
| EEG-DDM with keypoints | **$16.683 \pm 2.468$** | **$0.853 \pm 0.0151$** |

From the results depicted in Fig. 2 we observe that EEG-DDM performs better than other state of the art models (IC-UNet, ICLabel, Deep Separator) by demonstrating lower RRMSE and a higher CC. Furthermore, within the EEG-DDM framework, we demonstrate improvement in performance by utilising the keypoints and observing a lower RRMSE (**16.683** *±* **2.468** vs 22.281 *±* 1.278) and a higher correlation coefficient (**0.853** *±* **0.0151** vs 0.813 *±* 0.0156)

**Fig. 2:**
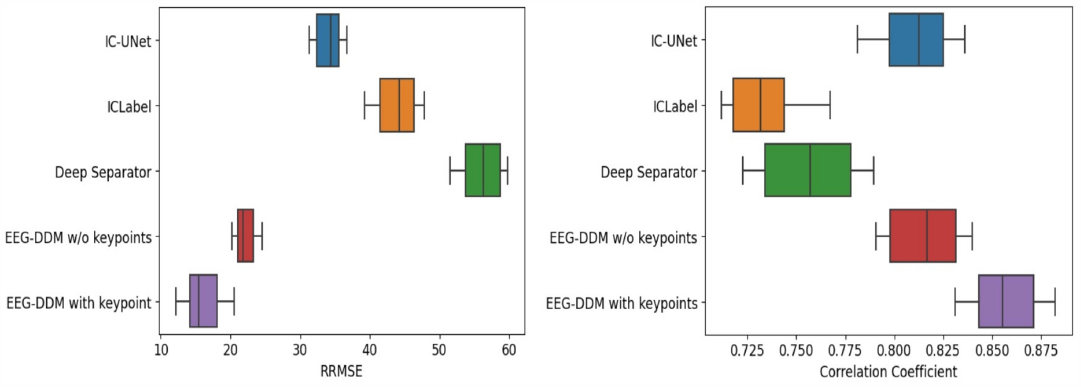
Performance of EEG-DDM with and without keypoints: The figure on the left represents the comparison of RRMSE with 3 different models (IC-UNet, ICLabel and Deep Separator) and EEG-DDM without keypoints and the plot in purple represents EEG-DDM RRMSE with keypoints. The figure on the right represents the same comparison with correlation coefficient as the metric

**Fig. 3:**
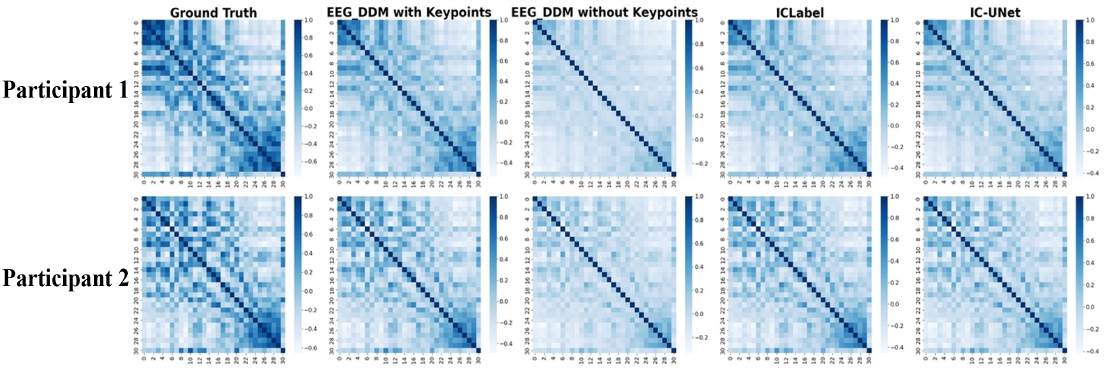
Visualisation of the Spatial Covariance matrices for 2 participants: The figure has 5 columns: the ground truth, EEG-DDM with keypoints, EEG-DDM without keypoints, ICLabel and IC-UNet where each column depicts the spatial covariance matrix estimated from the predicted clean EEG of that specific model.

### B. Preservation of Spatial Covariance with Keypoints

While denoising multichannel EEG, it is of importance to ensure that the denoised EEG upholds spatial covariance. The spatial covariance can be defined as *C* = *X* · *X*^*T*^ where X represents the clean EEG signal and in our case*X* ∈ **R**^31*×𝒯*^ .

We demonstrate the efficacy of keypoints by taking the Frobenius norm of the Euclidean distance between the ground truth EEG covariance matrix and the predicted covariance matrix.

From fig 5 we observe that the Euclidean distance of the covariance matrix estimated with the keypoints is smaller (mean: **8.895** and s.d **2.06**) as compared to the one estimated without the keypoints (mean; **12.537** and s.d **3.66**). From this, we can infer that keypoints improve the estimation of the noise variance and in turn the *t*. However do note that the model performs worse as compared to our baseline models without the keypoints.

#### C. Interpretability of Keypoint Contributions

As explained in section II-H, we visualise the mask ***M*** to explain the important key contributions for two different kinds of artifacts: ocular artifact and the muscular artifact. In fig 4, the mask ***M*** ∈ **R**^*C′×W*^ is plotted as a heatmap where the x axis represents samples in time and the y axis represents the feature respectively. In the case of ocular artifact, we observe that the feature indices corresponding to the y coordinate for both left and right eyes are highlighted in red representing their importance to the noise estimation. Furthermore, we observe a similar representation for the muscular artifact suggesting the noise source to be from the x coordinates of the neck, nose and both left and right ears. The change in x coordinates of all these features indicates a head tilt artifact. On the contrary, the blue regions indicate the regions of less interest to the model estimation and for both ocular and muscular artifacts, these regions are accurately identified as observed in fig 4

**Fig. 4:**
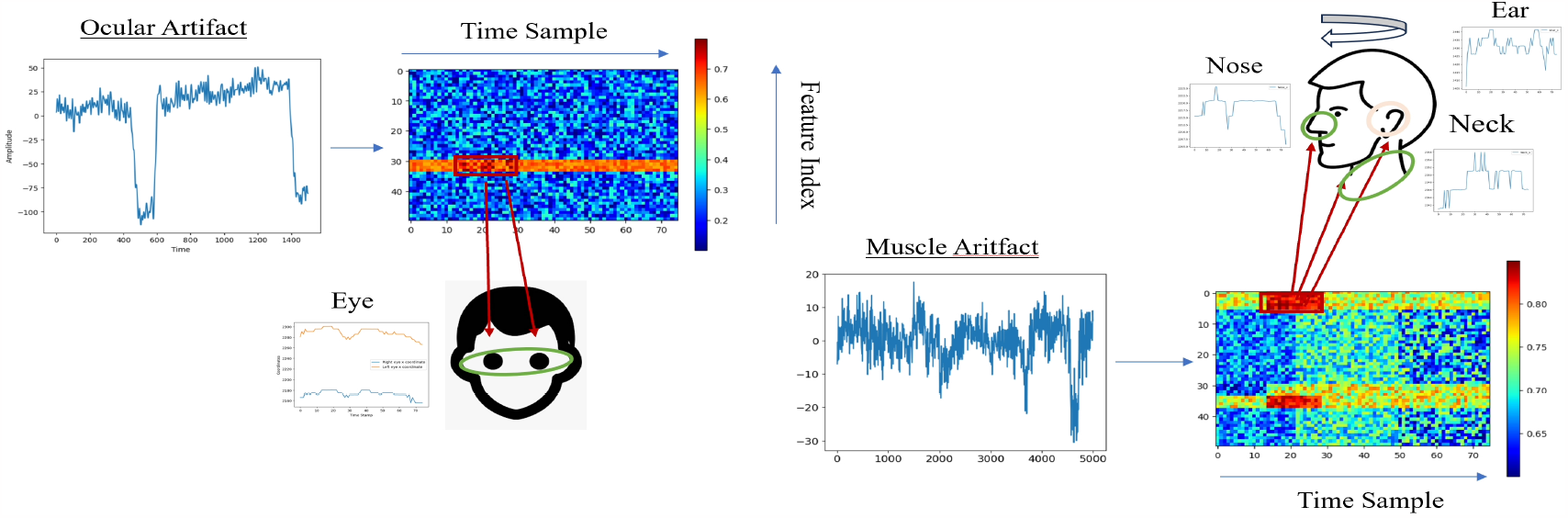
Visualisation of Saliency Maps generated from the Mask of the keypoint saliency model for Ocular and Muscular Noise: The y axis of the saliency map represents the feature index and the x axis represents the time dimension. The dark red regions of the saliency map represents high gradient values whereas the dark blue represents the lowest gradient values. In the figure on the left with the ocular noise of duration of 1500 samples, the high gradient values correspond to changes in the y coordinate of both left and right eye as depicted. In the figure on the right with muscular noise, the saliency maps depict a change in the x coordinates of nose, neck and both ear thus indicating the artifact may arise from a head tilt.

**Fig. 5:**
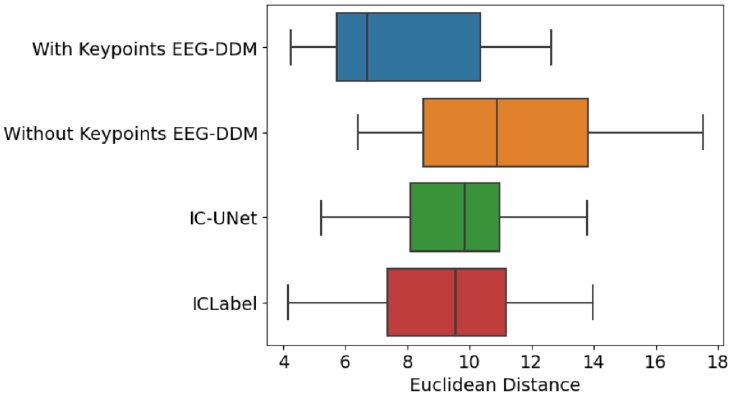
Comparison of Euclidean distance of spatial covariance across 15 participants

## IV. Conclusion

In this work, we address the challenge of removing artifacts from collected EEG signals in naturalistic settings for adults. In such environments, the signals are contaminated with non-stereotypical, high variance artifacts such as myogenic artifacts. We introduced a new modality - pose coordinates or body keypoints, estimated from a video source - to guide the denoising process that benefits, in particular, the removal of myogenic and ocular artifacts. To utilise these keypoints, we developed “EEG-DDM”, a score based diffusion model to denoise artifact contaminated EEG. Our results indicate that pose coordinates do help approximate the diffusion time *t* better than just utilising the artifact signal thus yielding superior performance not only in terms of RRMSE and correlation coefficient, but also a better approximation of realistic cross-channel spatial dependencies.

A limitation of our method is that multiple denoising steps are required to achieve a good denoising performance. This increases the time required to attain the clean EEG as compared to denoising models in prior works. One method of surpassing this limitation is by utilising an ODE solver which may be implemented in future work. Further, our model assumes the noise to follow an isotropic Gaussian which can be improved upon by assuming the noise variance evolution to be correlated across diffusion time steps, see for instance the approach in the recent work [21] that however does not consider denoising EEG artifacts.

## Acknowledgment

This research is supported by the RIE2025 Human Potential Programme Prenatal/Early Childhood Grant (H22P0M0002), administered by A*STAR

## Notes

### Competing Interest Statement

The authors have declared no competing interest.

## References

[1] B. L. Lee, “Understanding the usefulness of electroencephalography source localization,” Clinical and Experimental Pediatrics, vol. 66, no. 5, p. 210, 2023.

[2] X. Jiang, G.-B. Bian, and Z. Tian, “Removal of artifacts from eeg signals: a review,” Sensors, vol. 19, no. 5, p. 987, 2019.

[3] O. G. Lins, T. W. Picton, P. Berg, and M. Scherg, “Ocular artifacts in eeg and event-related potentials i: Scalp topography,” Brain topography, vol. 6, pp. 51–63, 1993.

[4] J. Ma, P. Tao, S. Bayram, and V. Svetnik, “Muscle artifacts in multichannel eeg: characteristics and reduction,” Clinical neurophysiology, vol. 123, no. 8, pp. 1676–1686, 2012.

[5] S. Leske and S. S. Dalal, “Reducing power line noise in eeg and meg data via spectrum interpolation,” Neuroimage, vol. 189, pp. 763–776, 2019.

[6] D. Langlois, S. Chartier, and D. Gosselin, “An introduction to independent component analysis: Infomax and fastica algorithms,” Tutorials in Quantitative Methods for Psychology, vol. 6, no. 1, pp. 31–38, 2010.

[7] C.-Y. Chang, S.-H. Hsu, L. Pion-Tonachini, and T.-P. Jung, “Evaluation of artifact subspace reconstruction for automatic eeg artifact removal,” in 2018 40th Annual International Conference of the IEEE Engineering in Medicine and Biology Society (EMBC). IEEE, 2018, pp. 1242–1245.

[8] D. Djuwari, D. K. Kumar, and M. Palaniswami, “Limitations of ica for artefact removal,” in 2005 IEEE Engineering in Medicine and Biology 27th Annual Conference. IEEE, 2006, pp. 4685–4688.

[9] E. Brophy, P. Redmond, A. Fleury, M. De Vos, G. Boylan, and T. Ward, “Denoising eeg signals for real-world bci applications using gans,” Frontiers in Neuroergonomics, vol. 2, p. 805573, 2022.

[10] J. Yin, A. Liu, C. Li, R. Qian, and X. Chen, “Frequency information enhanced deep eeg denoising network for ocular artifact removal,” IEEE Sensors Journal, vol. 22, no. 22, pp. 21 855–21 865, 2022.

[11] Z. Cao, G. Hidalgo, T. Simon, S.-E. Wei, and Y. Sheikh, “Openpose: realtime multi-person 2d pose estimation using part affinity fields. corr abs/1812.08008 (2018),” arXiv preprint arXiv:1812.08008, vol. 11, 2018.

[12] J. Piaget, “The construction of reality in the child,” Journal of Consulting Psychology, vol. 19, no. 1, p. 77, 1955.

[13] X. X. Tan and V. E. V. Leong, “A protocol for social interactive assessment of infant attention set-shifting between 12-24 months of age,” Available at SSRN 4461069, 2023.

[14] A. Delorme and S. Makeig, “Eeglab: an open source toolbox for analysis of single-trial eeg dynamics including independent component analysis,” Journal of neuroscience methods, vol. 134, no. 1, pp. 9–21, 2004.

[15] P. Dhariwal and A. Nichol, “Diffusion models beat gans on image synthesis,” Advances in neural information processing systems, vol. 34, pp. 8780–8794, 2021.

[16] Y. Song, J. Sohl-Dickstein, D. P. Kingma, A. Kumar, S. Ermon, and B. Poole, “Score-based generative modeling through stochastic differential equations,” arXiv preprint arXiv:2011.13456, 2020.

[17] H. Chung, E. S. Lee, and J. C. Ye, “Mr image denoising and superresolution using regularized reverse diffusion,” IEEE Transactions on Medical Imaging, vol. 42, no. 4, pp. 922–934, 2022.

[18] Q. Pan, W. Hu, and N. Chen, “Two birds with one stone: Series saliency for accurate and interpretable multivariate time series forecasting.” in IJCAI, 2021, pp. 2884–2891.

[19] D. P. Kingma and M. Welling, “Auto-encoding variational bayes,” arXiv preprint arXiv:1312.6114, 2013.

[20] P. Dabkowski and Y. Gal, “Real time image saliency for black box classifiers,” Advances in neural information processing systems, vol. 30, 2017.

[21] J. Vetter, J. H. Macke, and R. Gao, “Generating realistic neurophysiological time series with denoising diffusion probabilistic models,” bioRxiv, pp. 2023–08, 2023.

